# Prioritizing Non-coding Variants in Rice GWAS Loci with a Chromatin-Informed DNA Language Model

**DOI:** 10.64898/2026.07.24.740656

**Authors:** Josef Paolo A. Manlapaz, Anish M. S. Shrestha

## Abstract

Numerous genome-wide association studies in rice have identified loci associated with diverse agronomic traits. However, interpreting the regulatory and functional significance of these loci remains challenging because each locus often contains multiple variants in linkage disequilibrium, many of which lie in non-coding regions. Here, we present a method for prioritizing non-coding variants within an associated locus by integrating chromatin feature information predicted by a pretrained DNA language model fine-tuned on rice ChIP-seq and ATAC-seq datasets. We demonstrate the utility of our method through three case studies. In a post-GWAS analysis of heat tolerance, prioritized variants overlapped promoters of candidate genes previously identified through integrated GWAS and transcriptomic analyses, providing independent support for their potential regulatory roles. For the high-yield gene *DEP1*, promoter variant prioritization combined with *in silico* saturation mutagenesis identified a localized regulatory region enriched for high-impact mutations overlapping predicted transcription factor binding sites. For the drought-associated gene *OsHAK1*, the highest-ranked variant was predicted to be associated with chromatin features in a manner consistent with the reported co-occurrence of H2Bub with H3K4 methylation marks in plants. Overall, these results demonstrate the utility of our method for functionally informative prioritization of non-coding variants, facilitating the interpretation of GWAS loci and the identification of candidate regulatory variants in rice.

## 1. Introduction

Numerous genome-wide association studies (GWAS) have been conducted in rice (*Oryza sativa*) to identify single nucleotide polymorphisms (SNPs) associated with important agronomic traits [1]. Examples include GWAS on response to environmental stress such as salt tolerance [2], drought resistance [3], cold tolerance [4], and heat tolerance [5]; shape and size traits such as grain size [6–10] and leaf angle [11, 12]; metabolic traits [13, 14]; yield-related traits such as flowering time [15], plant compactness [16], awn length [17], and seed-setting rate [18].

Functional interpretation of GWAS output can be challenging. In rice, linkage disequilibrium can extend beyond 500 kb in some subpopulations [19], resulting in hundreds of highly correlated SNPs that exhibit similar association signals [20, 21]. Furthermore many of the associated SNPs tend to be located in non-coding regions, making it challenging to determine their functional relevance to the phenotype [22–24]. These issues necessitate post-GWAS analysis to prioritize the variants that provide biological insights into the mechanism underlying trait associations.

Functional interpretation of a GWAS locus can be improved by incorporating evidence from complementary experiments that provide biological context for associated variants in the locus. One important source of such information is ChIP-seq data which provide genome-wide maps of histone modifications. Several ChIP-seq datasets exist for rice [25–27]. Histone modifications are known to be associated with gene expression regulation during plant development, growth, and responses to environmental stimuli [28–30], and in fact have been shown to be predictive of gene expression levels [31, 32]. Importantly, studies have shown that histone modifications in regulatory regions can be predicted directly from the underlying DNA sequence [33]. This creates an opportunity to model the effect of non-coding variants in altering histone modification patterns, thereby providing a mechanistic basis for prioritizing candidate causal variants within associated loci.

Another related source of functional information is genome-wide chromatin accessibility profiles measured using ATAC-seq, which identifies open chromatin regions (OCRs) across the genome. Chromatin accessibility is closely linked to gene expression, as OCRs facilitate transcription factor (TF) binding and transcriptional activity [34, 35]. In non-coding regions, OCRs enable TF binding to cis-regulatory elements, thereby influencing gene expression [36, 37]. Similar to histone modifications, it has been shown that chromatin accessibility can be predicted from the underlying DNA sequence [38, 39]. This enables sequence-based models to predict the effects of non-coding variants on chromatin accessibility, providing functional evidence for prioritizing candidate causal variants within GWAS-associated loci in rice. Genome-wide ATAC-seq datasets have also been generated for rice [27], making similar sequence-based regulatory prediction approaches feasible in rice.

Several deep learning models have been developed to predict the regulatory effects of non-coding variants, enabling variant prioritization. One of the earliest models, DeepSEA [38], employed a convolutional neural network (CNN) to predict transcription factor binding, histone modifications, and chromatin accessibility directly from DNA sequence. By comparing predictions for the reference and alternative alleles, DeepSEA assigns regulatory effect scores to non-coding variants. Building on this framework, PlantDeepSEA [40] adapted the DeepSEA architecture for plant genomes, training species-specific models to predict chromatin accessibility across six plant species, including rice, and similarly uses predicted regulatory effects to prioritize variants. RiceVarMap [41] likewise employed a CNN trained on multi-tissue rice ATAC-seq data to predict chromatin accessibility and assign regulatory effect scores to variants. More recently, transformer-based architectures have emerged as an alternative to CNNs for regulatory effect prediction [42]. LOGO [43] employs a lightweight Bidirectional Encoder Representations from Transformers (ALBERT) architecture for predicting the regulatory effects of non-coding variants, while LOGOWheat [44] extends this approach to wheat genomes, demonstrating the potential of DNA language models for regulatory prediction in crop species.

More broadly, large pre-trained DNA language models such as DNABERT [45], Nucleotide Transformer [46], and DNABERT-2 [47] have been developed, leveraging BERT-based architectures pretrained on genomic sequences. Several of these models have also been further trained on plant genomic datasets, including Plant DNABERT [48] and AgroNT [49]. Pretraining on large-scale sequence datasets enables these models to capture contextual representations of DNA sequences, which can improve performance in downstream genomics tasks [50]. These models have been evaluated across a variety of applications, including promoter prediction, transcription factor binding site identification, and functional variant prediction, demonstrating improved performance in these areas [45–47]. Despite these advances, these models remain underutilized in the context of post-GWAS variant prioritization tasks for rice.

This study presents RiSPICE (Rice SNP Prioritization Integrating Chromatin Effects), which fine-tunes a large pre-trained DNA language model using rice genome-wide histone modification and chromatin accessibility profiles to predict the regulatory effects of non-coding variants. Similar to prior studies that applied this framework to plant genomes [40, 41, 44], RiSPICE assigns regulatory effect scores to non-coding variants based on predicted changes in chromatin features. These scores can be used to prioritize putative causal variants for downstream functional validation. The method is demonstrated through three case studies: (1) post-GWAS prioritization of variants within heat tolerance-associated loci, and promoter variant prioritization in (2) *DEP1*, a high-yield gene, and (3) *OsHAK1*, a drought-associated gene.

## 2. Methods

### 2.1. Overview of the Variant Prioritization Method

Our overall method is illustrated in Figure 1. It takes as input a set of trait-associated non-coding SNPs, each defined by a reference and an alternative allele. The method outputs two types of scores: the Per-Feature Score and the Overall Score. The Per-Feature Score represents the predicted effect of a SNP on an individual chromatin feature, whereas the Overall Score summarizes its predicted effect across all chromatin features. These outputs can be used to (1) rank SNPs based on the Overall Score, (2) examine the Per-Feature Scores to characterize the impact of each SNP across individual chromatin features, and (3) perform an *in silico* saturated mutagenesis analysis on a genomic region of interest, in which the Overall Score is computed for all possible single-nucleotide substitutions at every position within the region, enabling the identification of putative cis-regulatory elements and high-impact nucleotide positions.

**Figure 1:**
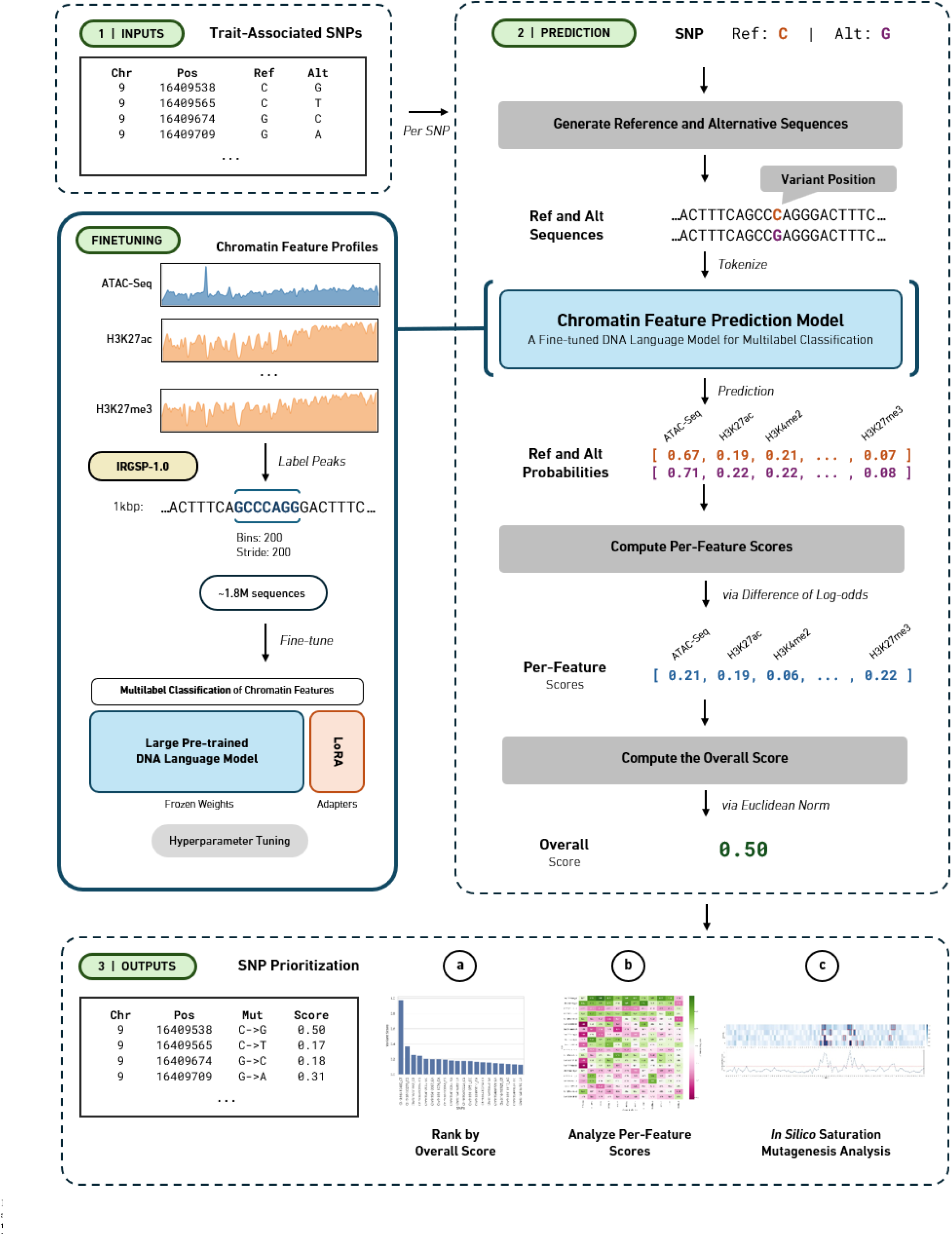
Overall RiSPICE variant prioritization method. **(1)** The method takes as input a set of trait-associated SNPs. **(2)** For each SNP, reference and alternative sequences are generated with the respective alleles centered at the variant position. These sequences are then provided as input to a chromatin feature prediction model fine-tuned to predict chromatin accessibility and histone modifications from an input DNA sequence. Using the predicted probabilities, Per-Feature Scores are computed as the difference in log-odds between the reference and alternative predictions, after which the Overall Score is computed as the Euclidean norm of the Per-Feature Scores. **(3)** The resulting Overall Score and Per-Feature Scores can then be used for variant prioritization by **(a)** ranking SNPs according to their Overall Score, **(b)** analyzing Per-Feature Scores across chromatin features, and **(c)** performing *in silico* saturation mutagenesis to identify high-impact sites within a genomic region.

At the core of our method is a chromatin feature prediction model, which is a DNA language model fine-tuned on rice ATAC-seq and ChIP-seq datasets to predict chromatin accessibility and histone modifications from an input DNA sequence. For each SNP, its Overall Score is computed following the approach used in prior studies [38, 40, 41, 43, 44]. We first generate reference and alternative sequences of length *L* bp, with the corresponding allele centered at the variant position in each sequence. These sequences are then used as inputs to the chromatin feature prediction model, which outputs reference and alternative probabilities representing the likelihood of each chromatin feature being present in the respective sequences. The Per-Feature Score is computed as the difference in log-odds between the reference and alternative probabilities. Finally, the Per-Feature Scores are aggregated across all chromatin features into the Overall Score using Euclidean normalization. Details of the Per-Feature Score and Overall Score computations, together with the methods used to assess their statistical significance, are presented in Section 2.5.

### 2.2 Training Data Collection

The *Oryza sativa* ssp. japonica cv. Nipponbare reference genome assembly (IRGSP-1.0) was retrieved from the NCBI Genome database (assembly accession `GCF_001433935.1`). ATAC-seq and ChIP-seq datasets for various histone modifications were primarily obtained from the RiceENCODE [27] database. Additional ChIP-seq datasets from Zheng et al. [25] were retrieved from the NCBI Gene Expression Omnibus (GEO; accession `GSE109618`). A summary of the datasets used in this study is provided in Table 1.

**Table 1.** Summary of Chromatin Features, Profiles, and Tissues used in the study.

| Features | Leaves | Roots | Panicles | Seedlings | SAM | Profile Count |
| --- | --- | --- | --- | --- | --- | --- |
| H3K23ac | • |  |  |  |  | 1 |
| H3K27ac | • | • |  |  |  | 2 |
| H3K27me3 | • | • |  |  | • | 3 |
| H3K36me3 | • | • |  | • |  | 3 |
| H3K4ac |  |  |  | • |  | 1 |
| H3K4me2 |  |  | • | • |  | 2 |
| H3K4me3 | • | • |  |  | • | 3 |
| H3K9ac |  | • | • | • |  | 3 |
| H3K9me1 |  |  |  | • |  | 1 |
| H4K12ac |  | • |  | • |  | 2 |
| H4K16ac | • |  |  |  |  | 1 |
| ATAC-Seq | • | • | • |  |  | 3 |
| **Totals** | **7** | **7** | **3** | **6** | **2** | **25** |

### 2.3. DNA Language Model Selection

We evaluated several published pre-trained DNA language models by assessing their ability to predict masked tokens in the Nipponbare reference genome. Based on its predictive performance and relatively low complexity, we selected DNABERT-2 [47] for downstream fine-tuning. We then fine-tuned three DNABERT-2 models using input sequence lengths of 500, 750, and 1000 bp, respectively, and selected the best-performing model for all subsequent analyses.

### 2.4. Fine-Tuning for Chromatin Feature Prediction

#### Generating Sequences

The Nipponbare reference genome was partitioned into non-overlapping bins of length *l*_core_, referred to as *core regions*. Each core region was extended by *n* bp upstream and downstream, resulting in sequences of length *L*, with the core region centered within each sequence. Sequences containing ambiguous nucleotides (`N`) were excluded. Following this procedure, we generated three sequence configurations with *L* = 500, 750, 1000 bp, corresponding to (*l*_core_, *n*) = (100, 200), (150, 300), (200, 400), respectively.

#### Identifying Peaks in ATAC-seq and ChIP-seq Profiles

For each ATAC-seq and ChIP-seq profile, peaks were called using `bdgpeakcall` and `bdgbroadcall` functions of `MACS3 v3.0.3` [51]. We initially used the default settings for these functions, including a `cutoff` of 5 (equivalent to a *p*-value of 1 *×* 10^*−*5^), a `min-length` of 200, and a `max-gap` of 30. These settings were treated as hyperparameters and iteratively tuned to improve model performance.

#### Dataset Construction

For each ChIP-seq and ATAC-seq peak file, we computed the overlap between the called peaks and every core region using `pybedtools`. The resulting coverage represented the proportion of each core region overlapped by a histone modification or chromatin accessibility peak. If at least 50% of a core region was covered by a peak, the corresponding sequence was assigned a label of 1 for that profile; otherwise, it was assigned a label of 0. Final labels for each chromatin feature were then determined by majority voting across all corresponding profiles.

The resulting dataset consisted of *L*-bp input sequences with a centered *l*_core_-bp core region, each assigned a binary label for every chromatin feature. The dataset was split by chromosome into training (chromosomes 1–5 and 8–12), validation (chromosome 7), and test (chromosome 6) sets. Chromosomes 6 and 7 were selected for the validation and test sets because their chromatin feature prevalence and sequence counts were closest to the genome-wide average across chromosomes. Each sequence was then tokenized using the published DNABERT-2 [47] tokenizer prior to model training and evaluation.

#### Training

We fine-tuned the pretrained DNABERT-2 model using Low-Rank Adaptation (LoRA) [52], where the pretrained model weights were frozen and only the LoRA adapter weights were optimized. After training, the adapter weights were merged with the base model to produce the final fine-tuned model. This procedure was applied independently to the three sequence configurations (*L* = 500, 750, and 1000 bp).

Binary cross-entropy (BCE) loss was used as the optimization objective during training. For each input sequence, the BCE loss was computed independently for each chromatin feature and summed across all *k* chromatin features:

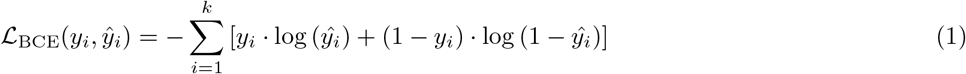

where *y*_*i*_ *∈* 0, 1 denotes the true label for the *i*-th chromatin feature, *ŷ*_*i*_ *∈* [0, 1] is the predicted probability for the *i*-th chromatin feature, and *k* is the total number of chromatin features.

The overall training loss was computed by averaging the per-sequence BCE loss across all *N* sequences:

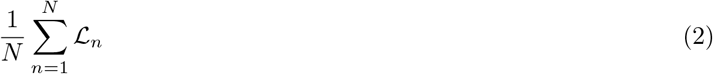

BCE loss was implemented using `BCEWithLogitsLoss` in `PyTorch`.

#### Evaluation

For each chromatin feature, AUROC and AUPRC were monitored on the validation set. We also computed Lift, defined as the ratio of AUPRC to prevalence, where prevalence is the proportion of genomic positions exhibiting the chromatin feature. Lift quantifies the model’s improvement over random selection in identifying positive instances and is particularly useful for evaluating performance on imbalanced datasets. Overall performance was obtained by macro-averaging.

Hyperparameters were iteratively tuned based on validation performance, including refinement of the `MACS3` peak-calling parameters and removal of chromatin profiles that negatively affected model performance. Once the final hyperparameter configuration was selected, the model was evaluated on the held-out test set.

### 2.5. Scoring and Statistical Significance

#### Computing the Per-Feature and Overall Scores

To compute the Per-Feature Scores for a SNP, we follow a similar approach used in prior studies [38, 40, 41]. For each chromatin feature *i* (e.g. H3K27ac), let *p*_*i*,alt_ and *p*_*i*,ref_ denote the predicted probabilities that chromatin feature *i* is present in the alternative and reference sequences, respectively. The Per-Feature Score, denoted by *δ*_*i*_, is computed as the difference in log-odds:

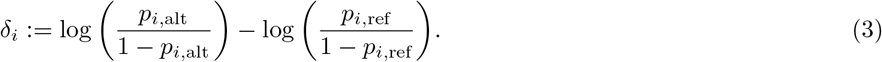

The Per-Feature Scores are aggregated across all chromatin features into an Overall Score using Euclidean normalization, following the approach of LOGOWheat [44]. The Overall Score, denoted as ∥*δ*∥, is computed as follows:

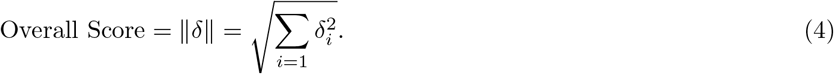

#### Statistical Significance

To construct empirical background distributions for determining the statistical significance of the Per-Feature Scores and Overall Score, we first randomly sampled 1 million SNPs genome-wide from the SNP-Seek [53] dataset. We included only non-coding variants, with the number of sampled variants from each chromosome proportional to the total number of variants present in that chromosome. The Per-Feature Scores and Overall Score were then computed for all the sampled SNPs. The resulting Overall Scores were used to construct an empirical background distribution from which significance thresholds were derived. In parallel, empirical background distributions were constructed separately for the Per-Feature Scores of each chromatin feature.

## 3. Results and Discussion

### 3.1. Our model reliably predicts chromatin features

The fine-tuned chromatin feature prediction models reliably predicted histone modifications and chromatin accessibility across all input sequence lengths (*L* = 500, 750, 1000 bp). Table 2 summarizes the performance of the three models, which achieved comparable results across all evaluation metrics. Among them, the 1000 bp model achieved the highest AUROC and AUPRC and was therefore selected for subsequent analyses.

**Table 2.** Overall performance of DNABERT-2 models trained using different sequence lengths.

|  | AUROC | AUPRC | Lift |
| --- | --- | --- | --- |
| 500 bp | 0.8884 | 0.6629 | 3.3785 |
| 750 bp | 0.8996 | 0.6761 | **3.6468** |
| 1000 bp | **0.9023** | **0.6946** | 3.5181 |

The performance of the selected 1000 bp model for each chromatin feature is shown in Table 3. The model achieved its highest predictive performance on H3K23ac, with an AUROC of 0.96, an AUPRC of 0.86, and a lift of 4.12. Although H3K27me3 and H3K4ac had relatively low AUPRC values (0.30 and 0.53, respectively), these histone modifications were also among the rarest in the fine-tuning dataset, with prevalences of 0.038 and 0.053. Consequently, the model achieved the highest lift values for H3K27me3 and H3K4ac (8.24 and 10.65, respectively), indicating that the model performed approximately 8.2*×* and 10.6*×* better than a random baseline.

**Table 3.** Predictive performance of the 1000 bp model for each chromatin feature.

| Chromatin Feature | AUROC | AUPRC | Lift |
| --- | --- | --- | --- |
| H3K23ac | **0.9629** | **0.8680** | 4.1265 |
| H3K27ac | 0.8514 | 0.7264 | 2.7872 |
| H3K27me3 | 0.8844 | 0.3083 | 8.2437 |
| H3K36me3 | 0.9252 | 0.7944 | 3.7114 |
| H3K4ac | 0.9413 | 0.5389 | **10.6540** |
| H3K4me2 | 0.9483 | 0.8470 | 3.5363 |
| H3K4me3 | 0.9117 | 0.7843 | 4.2940 |
| H3K9ac | 0.8795 | 0.7665 | 2.6858 |
| H3K9me1 | 0.8292 | 0.6869 | 2.3227 |
| H4K12ac | 0.9078 | 0.6581 | 5.5926 |
| H4K16ac | 0.8967 | 0.8257 | 2.1686 |
| ATAC-Seq | 0.8896 | 0.5309 | 5.6128 |
| **Average** | **0.9023** | **0.6946** | **3.5181** |

### 3.2. Prioritized Post-GWAS Heat Tolerance Variants are Consistent with Candidate Genes Identified by Integrated Transcriptomic Analysis

Recently Li et al. [5] conducted a GWAS on heat tolerance in rice using 620 rice accessions. Additionally, they integrated the GWAS results with transcriptomic data and identified 5 candidate genes – *LOC_Os07g48830, LOC_Os07g48630, LOC_Os07g48710, LOC_Os07g48570*, and *LOC_Os07g48760* – among which *LOC_Os07g48710* was considered the primary candidate gene based on significant expression level difference between two accessions.

We replicated their GWAS; and similar to the original study, we identified a lead SNP at `Chr7:29,140,282`, and defined the associated locus as the 155 kb interval spanning `Chr7:29,067,638-29,223,510`. The p-values obtained from GWAS on this locus are visualized in Figure 2. Based on SNP-Seek [53] and Ensembl [54], this locus contains 1071 SNPs and 28 annotated genes respectively. Of these, 390 were found to be located within promoter regions, which we consider here as 2000 bp upstream of transcription start sites (TSS).

**Figure 2.**
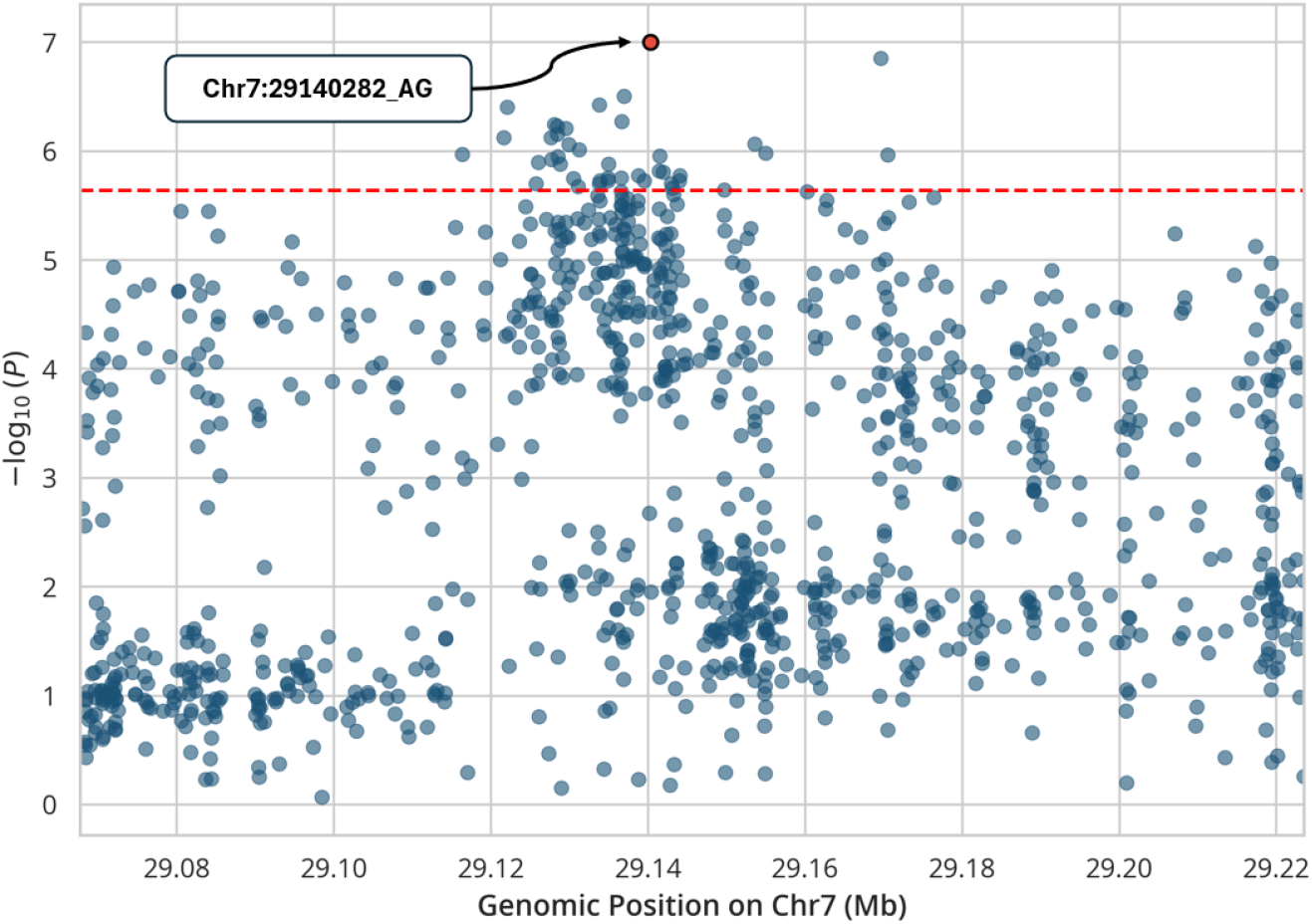
Regional association plot for the associated locus on chromosome 7, highlighting the lead SNP Chr7:29140282 AG. The red dashed line denotes the genome-wide significance threshold (*−* log_10_(*P*) = 5.64, corresponding to *P* = 2.29 *×* 10^*−*6^).

We computed the Overall and Per-Feature Scores of the 390 SNPs using our method. We identified 12 SNPs that met the 0.05 significance threshold. These SNPs are within the promoter regions of 8 RAP-DB [55] and MSU [56] genes, as summarized in Table 4. The top variant is located in the promoter of *LOC_Os07g48560* (*WOX11*), which is a gene known to be associated in abiotic stress in rice and other plants [57–59]. Further, two of these SNPs were located in the promoter region of the top candidate gene *LOC_Os07g48710* of the original study and two were located in the promoter region of another candidate gene *LOC_Os07g48570*. It is interesting that despite relying solely on chromatin feature information, our method identified variants consistent with those in the original transcriptomics-assisted post-GWAS analysis, providing independent support for the reported candidate genes.

**Table 4.**
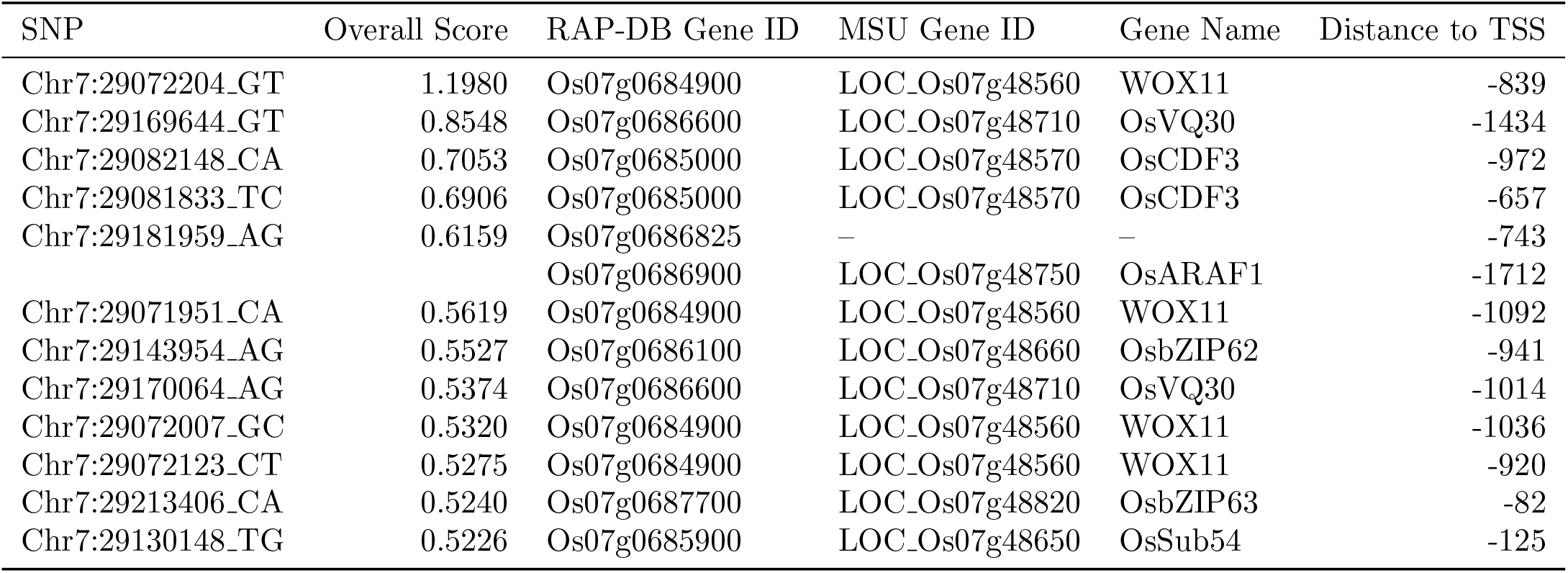
Promoter SNPs within the heat tolerance-associated interval (Chr7:29,067,638-29,223,510) that passed the significance threshold of 0.05, ranked by Overall Score.

Additionally, using RicePilaf [60], we identified clusters of genes in RCRN [61] that co-express with genes in the associated locus. The cluster containing *LOC_Os07g48560* was enriched for the Gene Ontology term cellular response to potassium ion starvation (GO:0051365). As potassium plays an important role in rice heat tolerance [62], this enrichment seems to be consistent with the heat tolerance phenotype associated with the GWAS locus. Co-expression network analysis also located *LOC_Os07g48570* (*OsCDF3*) in a cluster enriched for the Gene Ontology term temperature compensation of the circadian clock (GO:0010378), which has been associated with plant responses to abiotic stresses [63]. Notably, this gene was also identified as a candidate gene in the original study, and its promoter contains two of the significant SNPs listed in Table 4.

We next examined the Per-Feature Scores of the significant SNPs (Figure 3). The highest-ranked variant, `Chr7:29072204_GT`, was predicted to increase the likelihood of several chromatin features, including chromatin accessibility and multiple histone modifications, with H3K9me1 as the only feature predicted to decrease. In contrast, the second-ranked variant, `Chr7:29169644_GT`, showed the opposite pattern, with broadly reducing predicted chromatin accessibility and histone modification signals. Together, these predictions suggest that the two promoter variants may influence the local chromatin environment in distinct ways. Because chromatin accessibility and histone modifications are closely associated with gene regulation [28, 35], these variants represent plausible regulatory candidates for further functional validation.

**Figure 3.**
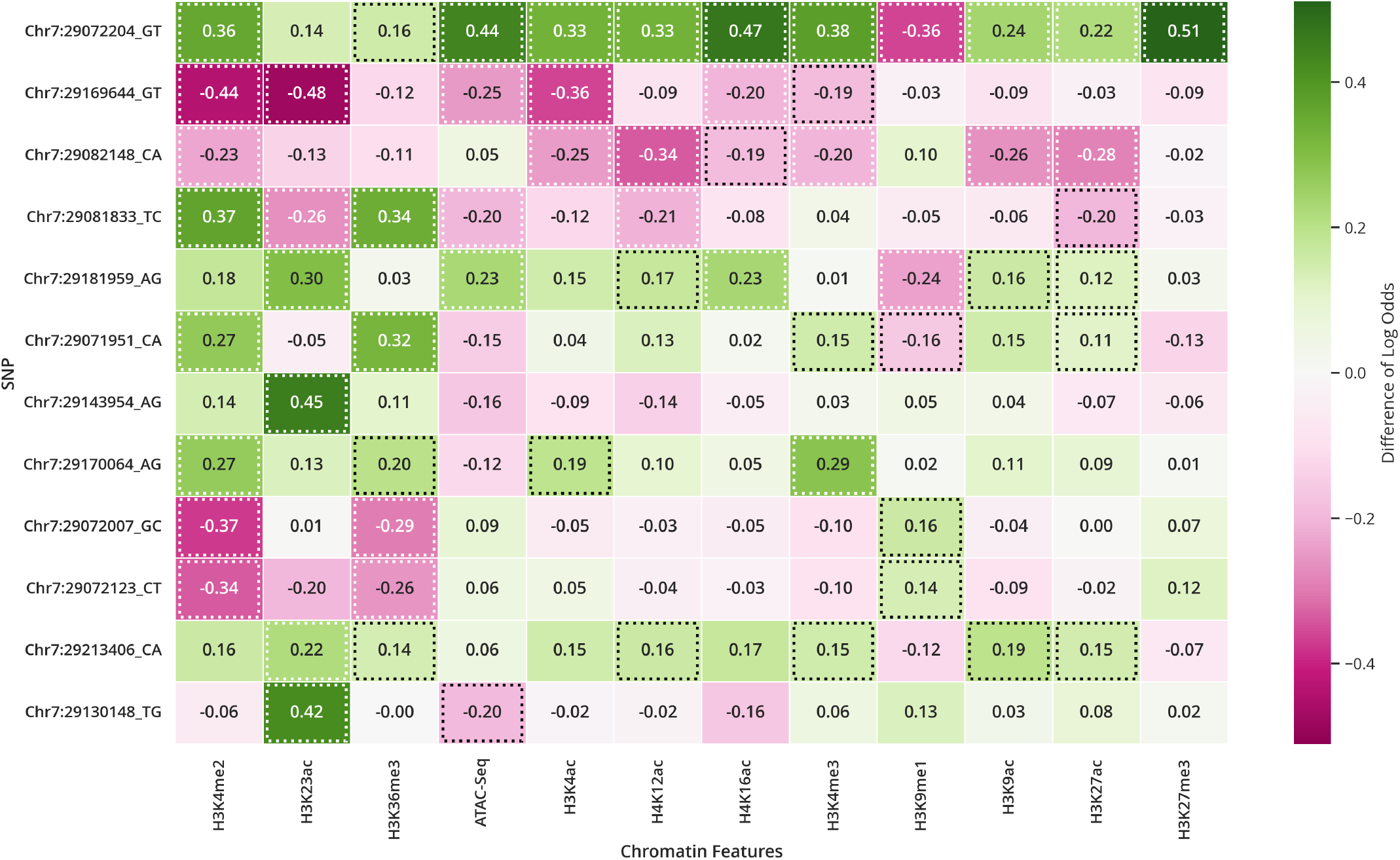
Heatmap of the Per-Feature Scores for the significant SNPs listed in Table 4. Green cells indicate that the alternate allele is predicted to increase the corresponding chromatin feature, whereas magenta cells indicate a predicted decrease. Cells outlined with dashed borders indicate Per-Feature Scores that exceeded the significance threshold of 0.05.

Considering the variant scores to be a test statistic for testing the lack of effect, we accounted for multiple testing within the locus by estimating the effective number of independent tests using the method of Li and Ji [64] and applied a Šidák correction [65], resulting in an adjusted significance threshold of 0.0126. Even under this more stringent criterion, the two highest-ranked variants remained significant: `Chr7:29072204_GT`, located in the promoter of *LOC_Os07g48560* (*WOX11*), and `Chr7:29169644_GT`, located in the promoter of *LOC_Os07g48710*, the primary candidate gene identified in the original study.

Collectively, these results demonstrate the utility of our framework for prioritizing and functionally characterizing non-coding variants in a post-GWAS context. By integrating predicted changes in chromatin accessibility and histone modifications, the framework provides biologically interpretable evidence that complements statistical association signals and facilitates the identification of candidate regulatory variants for downstream experimental validation.

### 3.3. Integrating Multiple Chromatin Features Refines Variant Prioritization in the *DEP1*

#### Promoter

The Dense and Erect Panicle 1 (*DEP1*) gene underlies the erect panicle phenotype, a high-yield breeding trait widely utilized in rice breeding [66, 67]. Because natural variation at the *DEP1* locus contributes to agronomic traits [68], and has previously been examined by PlantDeepSEA [40] and RiceVarMap [41], we re-examined the *DEP1* promoter region using our method.

We considered the 2000 bp region upstream of the *DEP1* TSS as the promoter region and retrieved 37 SNPs from this interval using the SNP-Seek database [53]. We computed the Overall and Per-Feature Scores for each variant using our method. Among them, `Chr9:16410405_CT` achieved the highest Overall Score and exceeded the multiple testing-adjusted significance threshold of 0.0141. As shown in Figure 4, this variant is clearly distinguished from the remaining promoter SNPs, highlighting its predicted impact.

**Figure 4.**
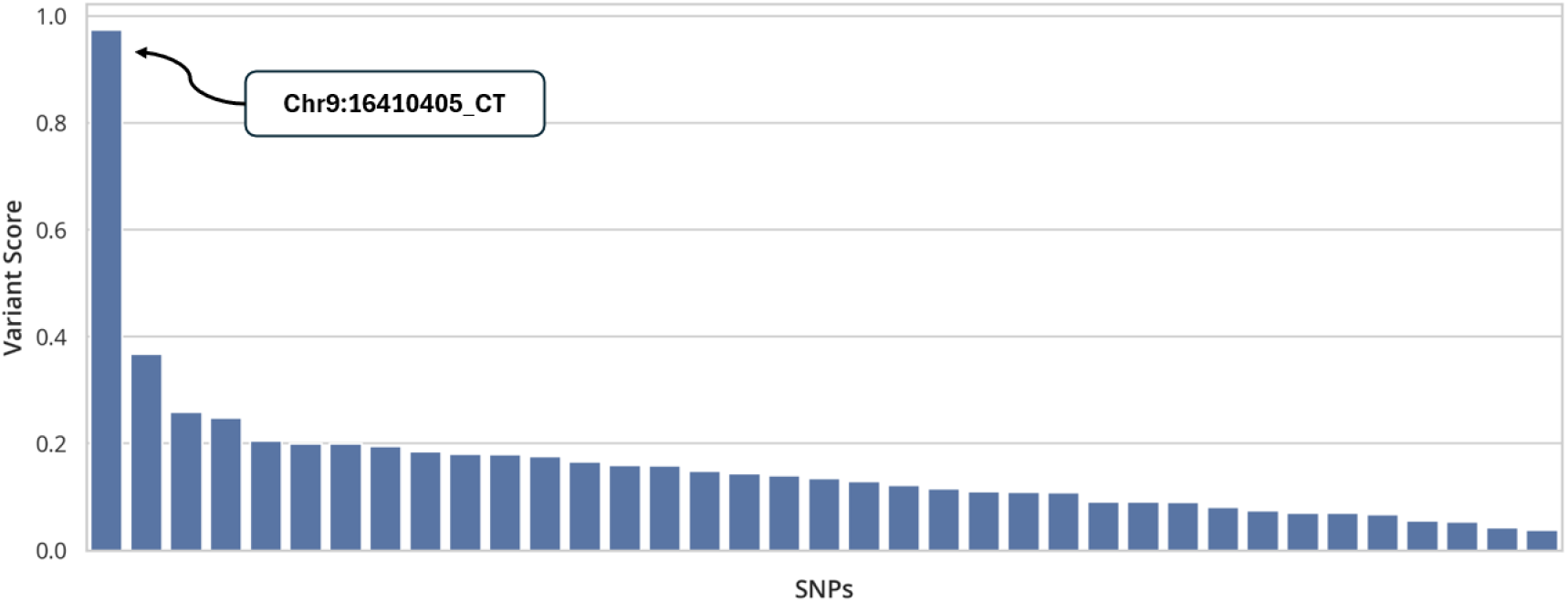
Ranked SNPs within the *DEP1* promoter region based on their Overall Scores. The highest-scoring variant, Chr9:16410405 CT, is highlighted.

We next examined the Per-Feature Scores of `Chr9:16410405_CT` (Figure 5). The variant was predicted to decrease the likelihood of most chromatin features, with the strongest reductions observed for H3K9ac, H3K27ac, H4K16ac, and H3K4me3. In contrast, it was predicted to increase the likelihood of chromatin accessibility and, to a lesser extent, H3K9me1.

**Figure 5.**
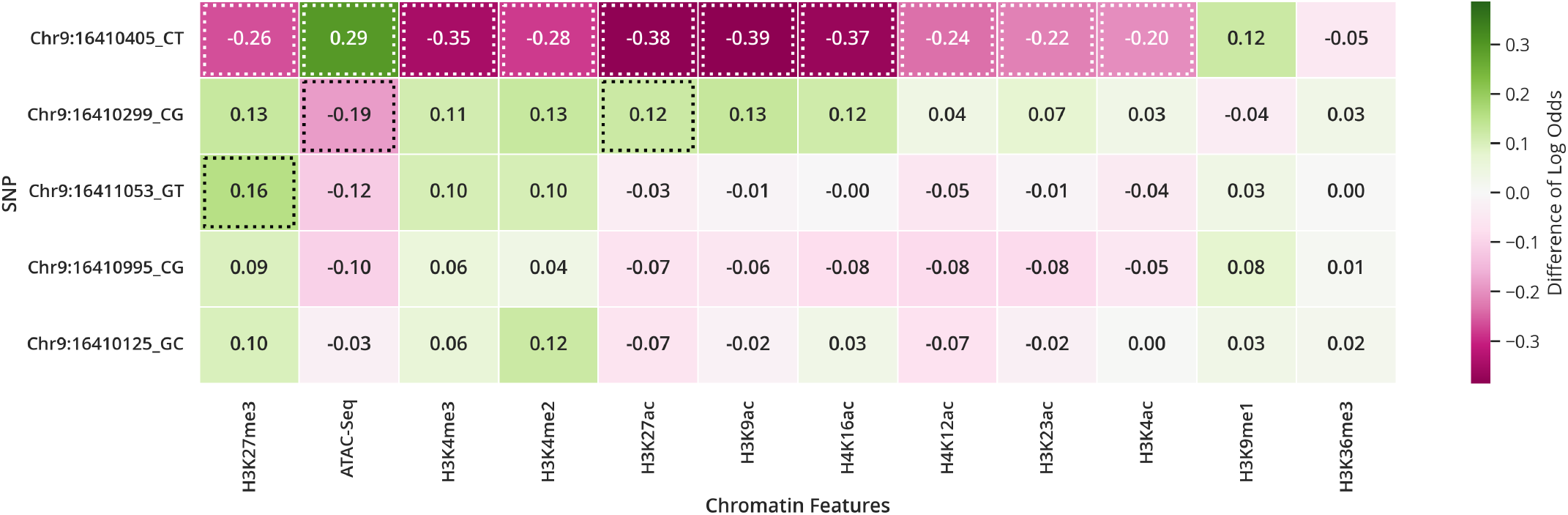
Heatmap of the Per-Feature Scores for the top five *DEP1* promoter variants ranked by Overall Score. Green cells indicate that the alternate allele is predicted to increase the corresponding chromatin feature, whereas magenta cells indicate a predicted decrease. Cells outlined with dashed borders indicate Per-Feature Scores that exceeded the significance threshold of 0.05.

Previous analyses using RiceVarMap [41] and PlantDeepSEA [40] identified a candidate regulatory variant at position `Chr9:16410299` in the *DEP1* promoter. Our analysis similarly ranked this variant highly, placing it second overall, and also predicted it to have a strong effect on chromatin accessibility (Figure 5). By integrating predicted effects across multiple chromatin features, our method instead ranked `Chr9:16410405_CT` as the highest-scoring promoter variant. The two variants are separated by 106 bp within the *DEP1* promoter, highlighting a localized region of predicted regulatory activity.

We performed *in silico* saturation mutagenesis analysis within a 100 bp region surrounding the top-ranked SNP `Chr9:16410405_CT`, in which we computed the effect of substituting each nucleotide in the region with all possible nucleotides (Figure 6). This analysis identified 20 significant substitution. Interestingly, the highest-scoring one was not the original SNP, but `Chr9:16410418_GT`. These significant substitutions were concentrated within two nearby intervals, Chr9:16,410,403–16,410,408 and Chr9:16,410,415–16,410,424, corresponding to the locations of the two highest-scoring variants. Further examination of these intervals showed that several high-impact substitutions overlapped predicted transcription factor binding sites (TFBSs) from PlantPAN [69], including sites associated with the AP2/ERF family transcription factor *Os07g0204000* and the bZIP transcription factor *HBF2* (*Os01g0813100*). The AP2/ERF family has been associated with rice growth and development [70], while *HBF2* has been implicated in flowering time regulation [71]. Together, the overlap between predicted TFBSs and high-impact mutations provides additional evidence that these variants may influence *DEP1* regulation.

**Figure 6.**
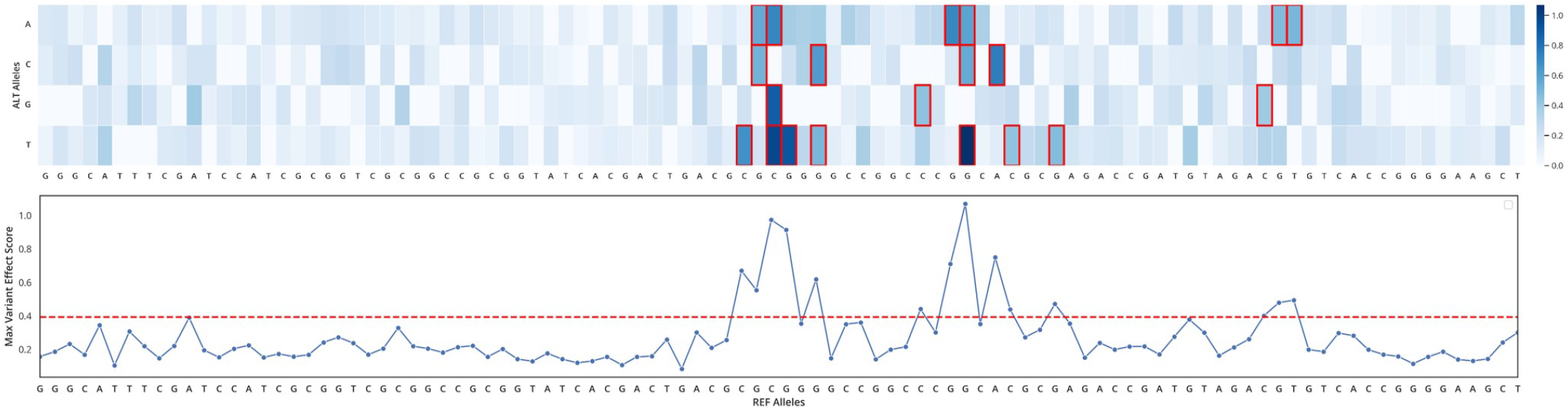
*In silico* saturation mutagenesis of the 100 bp region centered on the *DEP1* promoter SNP Chr9:16410405 CT. The heatmap shows the Overall Score for all possible nucleotide substitutions within the 100 bp region, where the x-axis represents the reference Nipponbare sequence and the y-axis represents the possible nucleotide substitutions at each position. Mutations exceeding the significance threshold of 0.1 are outlined in red. The line plot below summarizes the maximum Overall Score at each position, with the red dashed line indicating the significance threshold.

### 3.4. Predicted H3K4me2 Scores of *OsHAK1* Promoter Variants Are Consistent with H2Bub-Associated Regulation

It has been shown that histone H2B ubiquitination (H2Bub) plays an important role in the drought response of rice by regulating the expression of OsbZIP46 target genes [72, 73]. A recent study further showed that the transcription factor OsbZIP27 binds to the *OsHAK1* promoter, where it coordinates with *OsHUB1/2* to increase H2Bub and activate *OsHAK1* transcription under drought conditions [74]. H2Bub has also been associated with H3K4 methylation in plants, with H2Bub and H3K4me2/H3K4me3 reported to co-occur at transcriptionally regulated loci and act together in gene regulation [75–77]. H3K4me2 and H3K4me3 are among the chromatin features predicted by our model. Motivated by the established role of *OsHAK1* in drought tolerance [78] and evidence that H2Bub-mediated regulation at its promoter contributes to its expression, we examined variants within the *OsHAK1* promoter using our method.

We considered the 2000 bp region upstream of the *OsHAK1* TSS and retrieved 132 SNPs from this interval using the SNP-Seek database [53]. We computed the Overall and Per-Feature Scores for all variants. Of the 132 promoter SNPs analyzed, 5 exceeded the significance threshold of 0.05 and are summarized in Table 5.

**Table 5.** Promoter SNPs within the drought-associated gene *OsHAK1* that passed the significance threshold of 0.05, ranked by Overall Score.

| SNP | Overall Score | Distance to TSS |
| --- | --- | --- |
| Chr4:19884685_CA | 0.7795 | -1989 |
| Chr4:19884839_CT | 0.6722 | -1835 |
| Chr4:19885005_CA | 0.6116 | -1669 |
| Chr4:19886150_GA | 0.5350 | -524 |
| Chr4:19884767_CT | 0.5302 | -1907 |

We next examined the Per-Feature Scores of the five significant *OsHAK1* promoter variants (Figure 7). Among all chromatin features examined, the largest absolute Per-Feature Scores was observed for H3K4me2 across the prioritized variants. The highest-ranked variant, `Chr4:19884685_CA`, was predicted to decrease the likelihood of all chromatin features considered, with the largest reduction observed for H3K4me2, followed by H4K16ac, H3K9ac, and chromatin accessibility. In contrast, the third-ranked variant, `Chr4:19885005_CA`, showed the opposite pattern, with broadly increased predicted chromatin features, including H3K4me2, H3K4me3, and chromatin accessibility. The remaining significant variants were also predicted to predominantly reduce chromatin feature likelihoods. As H2Bub has been reported to act in concert with H3K4 methylation in plants [75], the consistently large H3K4me2 Per-Feature Scores provide biological context for the prioritized variants.

**Figure 7.**
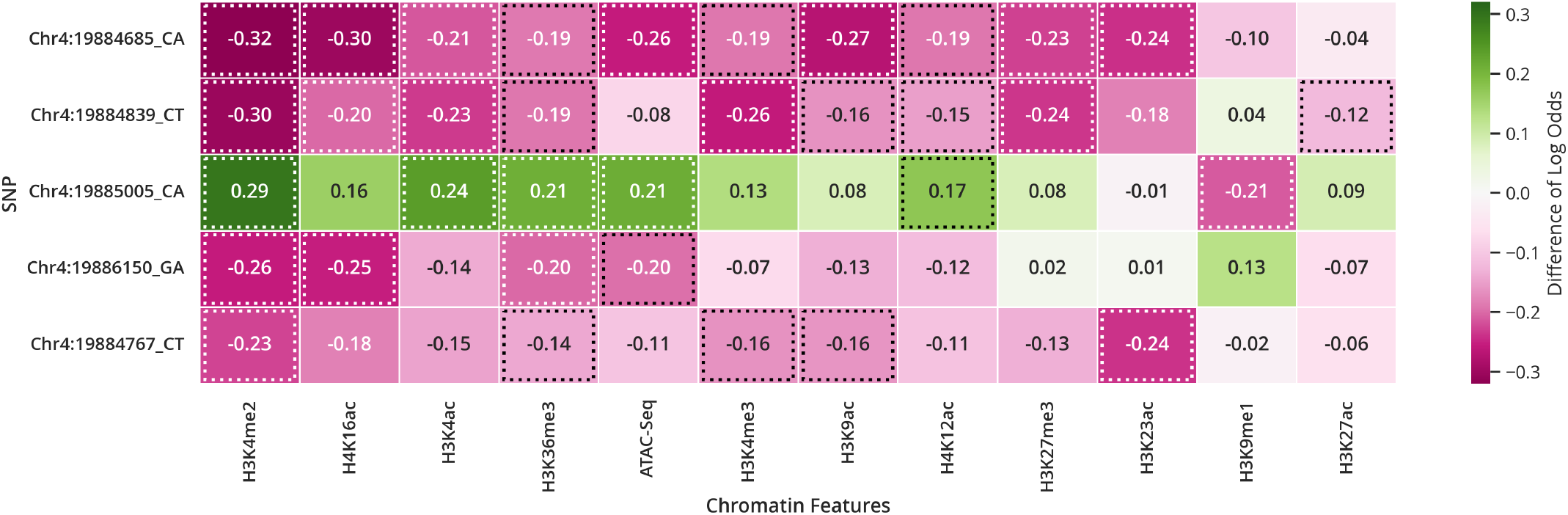
Heatmap of the Per-Feature Scores for the significant SNPs listed in Table 5. Green cells indicate that the alternate allele is predicted to increase the corresponding chromatin feature, whereas magenta cells indicate a predicted decrease. Cells outlined with dashed borders indicate Per-Feature Scores that exceeded the significance threshold of 0.05.

To further examine the sequence context surrounding the prioritized *OsHAK1* variant, we performed *in silico* saturation mutagenesis across a 100 bp region centered on `Chr4:19884685_CA`, in which we computed the effect of substituting each nucleotide in the region with all possible nucleotides (Figure 8). We identified significant substitutions at 44 positions, indicating that the surrounding sequence contains multiple putative regulatory sites. Many of these high-impact substitutions overlapped predicted TFBSs from PlantPAN [69], including sites for the ABA-responsive transcription factor *OsBZ8* [79]. Given the central role of ABA signaling in drought responses [80], these findings provide additional support for the regulatory relevance of this interval within the *OsHAK1* promoter.

**Figure 8.**
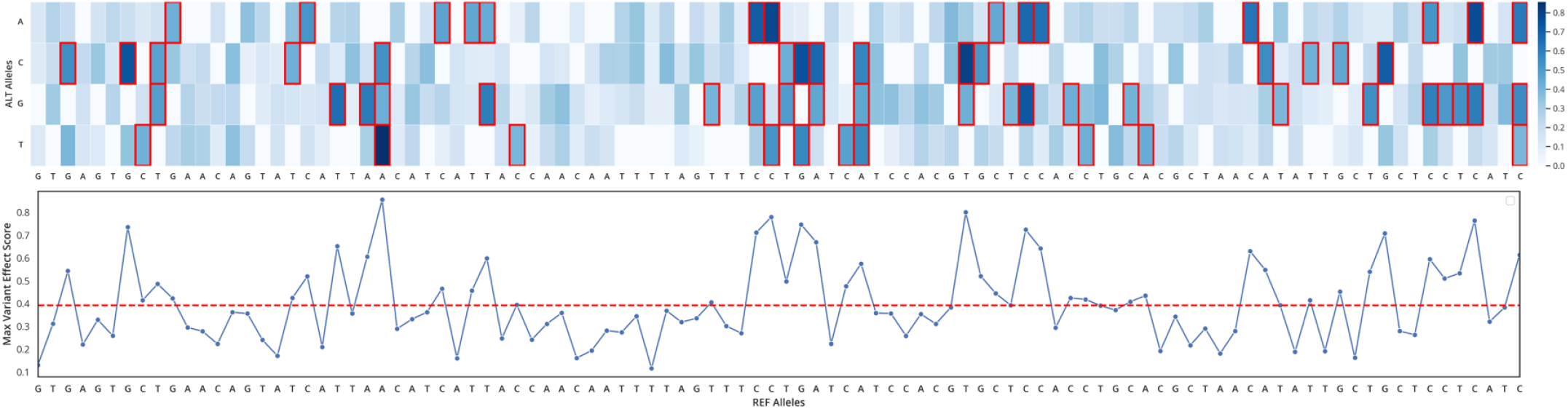
*In silico* saturation mutagenesis of the 100 bp region centered on the *OsHAK1* promoter SNP Chr4:19884685 CA. The heatmap shows the Overall Score for all possible nucleotide substitutions within the 100 bp region, where the x-axis represents the reference Nipponbare sequence and the y-axis represents the possible nucleotide substitutions at each position. Mutations exceeding the significance threshold of 0.1 are outlined in red. The line plot below summarizes the maximum Overall Score at each position, with the red dashed line indicating the significance threshold.

## 4. Conclusion

In this study, we demonstrated how RiSPICE leverages a DNA language model to prioritize variants within loci associated with agronomic traits in rice. We fine-tuned DNABERT-2 on publicly available rice histone modification and chromatin accessibility data and showed that it reliably predicts chromatin features directly from sequences. We showed through three case studies – a heat tolerance locus, the promoter region of *DEP1*, and a drought tolerance associated gene *OsHAK1* – that our model is able to prioritize variants in a biologically meaningful way. Compared to existing models [40, 41], our approach supports a broader set of chromatin features specific to rice, allowing for a more comprehensive analysis of the chromatin landscape.

A primary limitation of this study is the limited availability of public rice datasets. The current model was trained using chromatin profiles collected from RiceENCODE [27] and related studies [25], which restricted the analysis to 11 histone modifications and chromatin accessibility (ATAC-seq). Although these features provide a useful baseline for modeling the regulatory effects of non-coding variants, a more complete view of the regulatory landscape would require additional epigenomic datasets. Future versions of the pipeline could incorporate other modalities such as DNA methylation, additional histone marks, and transcription factor binding site profiles.

Another limitation is the lack of comprehensive tissue-specific chromatin feature profiling in rice. Because of this, the present study used a generalized modeling approach in which chromatin features were aggregated across tissues, rather than predicting tissue-specific chromatin states. As more high-quality tissue-specific ChIP-seq and ATAC-seq datasets become available, the framework could be extended to model chromatin features on a per-tissue basis. This would allow more context-specific analyses, including tissue-specific variant prioritization, chromatin landscape analysis, and *in silico* saturated mutagenesis across different tissues or developmental stages.

Finally, the model was fine-tuned using datasets mapped to the *Oryza sativa* cv. Nipponbare reference genome. While this makes the model directly relevant for analyses based on the Nipponbare genome, it may limit its applicability to other rice cultivars with different sequence backgrounds. Nevertheless, the overall pipeline is transferable, and similar models could be trained for other rice genomes as comparable epigenomic datasets become available.

## Data Availability

The datasets used in this study were obtained from publicly available resources as described in the Methods section. The RiSPICE repository (https://github.com/bioinfodlsu/rispice) provides links to the fine-tuned LoRA adapters and the original pretrained DNABERT-2 model used in this study.

## Availability of Source Code and Requirements

- Project Name: RiSPICE
- Project Home Page: https://github.com/bioinfodlsu/rispice
- Operating System(s): Platform Independent
- Programming Language: `Python 3.10.13`
- Other Requirements: `PyTorch`; Hugging Face Transformers; `PEFT`; `NumPy`; `pandas`
- License: MIT License
- Bio.tools ID: `biotools:rispice`

